# Identification of *Avramr1* from *Phytophthora infestans* using long read and cDNA pathogen-enrichment sequencing (PenSeq)

**DOI:** 10.1101/2020.05.14.095158

**Authors:** Xiao Lin, Tianqiao Song, Sebastian Fairhead, Kamil Witek, Agathe Jouet, Florian Jupe, Agnieszka I. Witek, Hari S. Karki, Vivianne G. A. A. Vleeshouwers, Ingo Hein, Jonathan D. G. Jones

## Abstract

- Potato late blight, caused by the oomycete pathogen *Phytophthora infestans*, significantly hampers potato production. Recently, a new *Resistance to Phytophthora infestans* (*Rpi*) gene, *Rpi-amr1*, was cloned from a wild *Solanum* species, *Solanum americanum*. Identification of the corresponding recognized effector (*Avirulence*, or *Avr)* genes from *P. infestans* is key to elucidating their naturally occurring sequence variation, which in turn informs the potential durability of the cognate late blight resistance.
- To identify the *P. infestans* effector recognized by *Rpi-amr1*, we screened available effector libraries and used long read and cDNA pathogen-enrichment sequencing (PenSeq) on four *P. infestans* isolates to explore the untested effectors.
- By using SMRT and cDNA PenSeq, we identified 47 highly expressed effectors from *P. infestans*, including PITG_07569 which triggers a highly specific cell death response when transiently co-expressed with *Rpi-amr1* in *Nicotiana benthamiana*, suggesting that *PITG_07569* is *Avramr1*.
- Here we demonstrate that long read and cDNA PenSeq enables the identification of full-length RxLR effector families, and their expression profile. This study has revealed key insights into the evolution and polymorphism of a complex RxLR effector family that is associated with the recognition by *Rpi-amr1*.

## Introduction

Potato late blight, caused by the hemi-biotrophic oomycete pathogen *Phytophthora infestans*, triggered the Irish and European famine in the late 1840s, and still causes severe losses to world potato production.

To reduce losses, breeders sought resistance genes in wild relatives of potato. Early in the 20^th^ century, *Solanum demissum*, a highly resistant hexaploid (2n=72) wild potato was found to be a useful source of Resistance to *P. infestans (Rpi)* genes (Salaman, 1937). Since then, many resistance traits have been transferred to cultivated potatoes by introgression breeding (Toxopeus, 1956), and many *Rpi* genes have been cloned from wild potatoes, e.g. *R1*, *R3a*, *R8*, *Rpi-blb1* and *Rpi-vnt1* (Ballvora *et al.*, 2002; van der Vossen *et al.*, 2003; Huang *et al.*, 2005; Foster *et al.*, 2009; Pel *et al.*, 2009; Vossen *et al.*, 2016). Unlike wild potatoes, *Solanum nigrum* and *Solanum americanum* have been reported to be non-hosts for *P. infestans* (Colon *et al.*, 1993). Two *Rpi* genes encoding NLR proteins, *Rpi-amr3* and *Rpi-amr1*, were cloned from *S. americanum* and confer late blight resistance in potato (Witek *et al.*, 2016; 2020).

Identification of the recognized effectors for *Rpi-amr3* and *Rpi-amr1* would open the way to investigate their virulence function and distribution in *P. infestans* populations. Moreover, it could also help to diagnose *Rpi* gene repertoires in resistant plants, and individually confirm their activity in genetically modified potatoes carrying multiple *Rpi* genes. In oomycetes, all the cloned Avr proteins contain a signal peptide and RxLR motif (Rehmany *et al.*, 2005), and the genomic sequencing of *P. infestans* revealed 563 RxLR effectors in the T30-4 reference genome (Haas *et al.*, 2009). This enabled a high-throughput effectoromics approach for functional screening of the candidate effectors in plants (Vleeshouwers *et al.*, 2008; 2011), and many *Avr* genes were identified by this approach, including *Avrblb1*, *Avrblb2* and *Avrvnt1* (Vleeshouwers *et al.*, 2008; Oh *et al.*, 2009; Pel, 2010).

However, available RxLR effector libraries do not contain recombinant clones of all *P. infestans* RxLR effectors, the effector candidates were defined on the basis of expression profile, motif analysis and distribution between *P. infestans* races (Vleeshouwers *et al.*, 2008; Oh *et al.*, 2009; Haas *et al.*, 2009). In total, ~300/563 RxLR effectors were previously cloned into expression vectors for functional screening (Rietman, 2011).

To further explore the diversity of RxLR effectors from *P. infestans*, a pathogen enrichment sequencing (PenSeq) approach was adopted to study allelic variation of RxLR effectors and population genomics of oomycetes. A bait library of RxLR effectors and some other pathogen-related genes was synthesized and used for enrichment prior to sequencing (Jouet *et al.*, 2018; Thilliez *et al.*, 2018). However, the previous PenSeq analyses used Illumina reads and genomic DNA (gDNA), making it difficult to differentiate individual effector alleles and closely related paralogs, or to find out which effectors are expressed.

Here, to identify the recognized effector of the newly-cloned Rpi-amr1 protein from *S. americanum* (Witek *et al.*, 2020), we screened all currently available RxLR effectors for recognition but without success. Therefore, we used PenSeq with long read (PacBio) and cDNA methods, and extended the list of candidate effectors that could be screened. Amongst these additional candidate RXLR genes, we identified *Avramr1* and defined orthologs and paralogs from four different isolates of *P. infestans*.

## Materials and Methods

### Sample preparation

To collect the mycelium of *Phytophthora infestans* for DNA extraction, *P. infestans* strains were grown on RSA solid medium for 7 days and then moved to Plich liquid media for 14 days. Mycelia were washed and harvested, freeze-dried using a vacuum pump and stored at −80°C until DNA or RNA extraction.

To collect the infection samples or zoospores for RNA extraction, *P. infestans* strains were cultured for 10 days on RSA medium. Grown mycelia were covered with cold (4 °C) sterile water and then incubated at 4-6 °C for 2-3h. The concentration of the inoculum was adjusted to about 50,000 zoospores/mL and 10 μL drops of inoculum were placed on the detached leaves of potato plants. Detached leaf assays (DLA) were incubated at 20 °C in high humidity for a required time post inoculation. Leaf discs of the infection area were collected and stored at −80°C until DNA or RNA extraction.

### DNA and RNA extraction

DNA was extracted using phenol/chloroform. *P. infestans* mycelium samples or infected leaf discs were ground into powder in liquid nitrogen. Ground material was resuspended in 500 μl of Shorty buffer (20% 1M Tris HCl pH 9, 20% 2M LiCl, 5% 0.5M EDTA, 10% SDS 10%, 45% dH2O) and one volume of phenol: chloroform: isoamyl alcohol (25: 24: 1) was added. The upper aqueous phase containing DNA was mixed with one volume of 100% ice-cold isopropanol to precipitate DNA. The pellet was washed twice using 70% ethanol, heated at 70°C for 2-5 minutes to completely remove ethanol and resuspended in sterile water. Resuspended DNA was then heated at 65°C for 20 minutes to inactivate DNases before RNase treatment was performed (2 μl of 10 mg/ml^−1^ RNase A, 37°C, 1h) and RNase A removed by chlorophorm precipitation. Genomic DNA was resuspended in water and sheared into 3-5 kb fragments using the S220 Focused-ultrasonicator (Covaris Inc., MA, USA).

RNA samples were extracted with Direct-zol™ RNA MiniPrep kit (Zymo Research, Tustin, CA, USA) according to the manufacturer’s instructions.

### PacBio and Illumina PenSeq capture

PacBio library was constructed with DNA samples from the mycelium of four *P. infestans* strains, EU_13_A2 (2006_3928A), EC1_A1 (EC1_3626), EU_6_A1 (2006_3920A) and US23. The library construction and target DNA sequence capture were performed according to (Witek *et al.*, 2016), with minor modifications. Qubit Fluorometer (ThermoFisher, Dubuque, IA, USA) was used to quantify the barcoded DNA library from each isolate. Equimolar amounts of DNA from the four individually barcoded samples were pooled to obtain 250 ng of total DNA and then subjected to sequence capture. A 10x excess of non-adaptor-ligated *P. infestans* DNA at about 500-1,000 bp was added for the hybridization. The final mixture of the amplicons of the captured library was further size selected by SageELF electrophoresis system (Sage Science, MA, USA) according to the instructions of the manufacturer.

Illumina library was constructed with RNA samples from zoospores of the four *P. infestans* strains, from the corresponding infected leaf discs harvested at 12 hours post infection (hpi), 1, 2 and 3 days post infection (dpi), and from mycelium of EU_13_A2. An Illumina library for each sample was constructed with KAPA mRNA HyperPrep Kit for Illumina® Platforms (KR1352 – v5.17) following the manufacturer’s instructions. mRNA was fragmented to 300-400 bp. The barcoded libraries were mixed together at a ratio of 16: 8: 4: 1: 1 for 12 hpi, 1 dpi, 2 dpi, 3 dpi, zoospores and mycelium samples, respectively.

Both types of libraries were subjected to sequence capture using the bait library as described previously (Jouet *et al*., 2018, Thilliez *et al.*, 2018). Before and after sequence capture, qPCR was performed on Bio-Rad CFX96 real-time detection system with an input of 1 ng DNA to assess the efficiency of capture.

### Sequencing

PacBio PenSeq libraries were sequenced at the Earlham Institute (Norwich, UK) using Sequel platform. Illumina PenSeq cDNA libraries were sequenced at Novogene (Hong Kong, China) using HiSeq, PE250.

### gDNA PenSeq assembly

PacBio raw reads were processed as described in (Witek et al., 2016) to generate ROI reads and demultiplexed using custom script (Van de Weyer et al., 2019). Demultiplexed ROI were assembled using Geneious R8 (http://www.geneious.com/) using settings as in (Witek et al., 2016).

### Analysis of cDNA PenSeq

All RxLR effectors from the *P. infestans* reference genome T30-4 were used to generate an artificial “RxLRome” contig, where RxLR effectors’ sequences were separated by stretches of 500 “Ns”. The contig also contained nine non-RxLR control genes (Jouet et al., 2018). The cDNA PenSeq reads from all treatments were mapped to the T30-4 RxLRome, and the expression analyses were performed and visualized using Geneious R10 (Kearse et al., 2012).

### New candidate RxLR effectors

For the previously untested RxLR effectors, we first selected the effectors showing differential expression at different stages and ranked them based on the raw transcript counts. Next, local alignment searches (BLAST) were performed against the 563 predicted RxLR effectors (Haas *et al.*, 2009) to remove the previously tested effectors. This analysis revealed 47 candidate RxLR effectors which were not included in previous functional study. The 47 RxLR effectors were synthesized by Twist Bioscience (San Francisco, CA, USA). The signal peptides were removed, the sequences were domesticated for Golden Gate cloning, and overhangs containing BsaI restriction sites were added to both ends of all effector sequences.

All the effectors were cloned into vector pICSL86977 (TSL SynBio) with CaMV 35S promoter and OCS terminator. The constructs were transformed to *Agrobacterium* strain GV3101 for agro-infiltration.

### Cell death assay

Transient expression of RxLR effectors and *Rpi-amr1* in *Nicotiana benthamiana* was performed as described previously (Bos *et al.*, 2006). *Agrobacterium* was infiltrated at OD600=1, and each effector was co-infiltrated with *Rpi-amr1-2273* (Witek *et al.*, 2020). The cell death phenotype was observed at 4 dpi.

### Data availability

Raw PacBio and cDNA PenSeq read sequences have been deposited in the Sequence Read Archive (SRA) under BioProject IDs PRJNA623167 and PRJNA598824.

## Results

### Available recombinant RxLR effector libraries do not contain *Avramr1*

To identify *Avramr1*, we tested 278 available RxLR effectors (Table S1) by co-expressing them with *Rpi-amr1-2273 i*n *N. benthamiana* (Rietman, 2011; Witek *et al.*, 2020). However, no effector activated *Rpi-amr1*-dependent HR, so we concluded that *Avramr1* was absent from the available RxLR effector libraries. Notably, *Avr8* was not originally included in the core effector selection, because *Avr8* expression goes up earlier then 2 dpi (Jo, 2013), showing that the criteria adopted to define core effectors did not reveal all recognized effectors.

To find *Avramr1*, we proposed three hypotheses: 1) *Avramr1* is an RxLR effector but it is not present in the assembled version of *P. infestans* T30-4 reference genome 2) *Avramr1* is an RxLR effector but it was not yet tested in previous functional studies/libraries; 3) *Avramr1* is not a typical RxLR effector. To address hypothesis 1, we performed PacBio PenSeq to sequence the effector alleles in the four diverse *P. infestans* isolates, EU_13_A2 (2006_3928A), EC1_A1 (EC1_3626), EU_6_A1 (2006_3920A) and US23, all of them avirulent on potato plants carrying *Rpi-amr1*, it indicates they all carry the recognized effector. To address hypothesis 2, we performed cDNA PenSeq to try to identify other RxLR effectors that are expressed during infection but not reported or defined in previous functional studies.

### PacBio PenSeq of four *P. infestans* isolates EU_13_A2, EC1_A1, EU_6_A1 and US23

PacBio gDNA PenSeq was performed on four *P. infestans* isolates of genotypes EU_13_A2, EC1_A1, EU_6_A1 and US23 (Fig. 1a). To evaluate the enrichment efficiency, qPCR was performed with the DNA pre-and post-capture. In general, the targeted genes of different length were well enriched at Concentration x time (Cot) value <20, while the untargeted genes were almost undetectable, with Cot value >27 (Peterson *et al.*, 2002) (Fig. S1). Furthermore, we found that the capture efficiency was increased by including a 10-fold molar excess of non-adaptor-ligated fragmented *P. infestans* DNA (500-1,000 bp) in the reannealing reaction, to reduce the extent to which sequences were recovered due to concatenation of transposon-containing sequences adjacent to *RxLR* genes. After sequence capture, enrichment of most effector genes was more efficient when non-adaptor-ligated *P. infestans* DNA was included (Fig. S1).

**Fig. 1.**
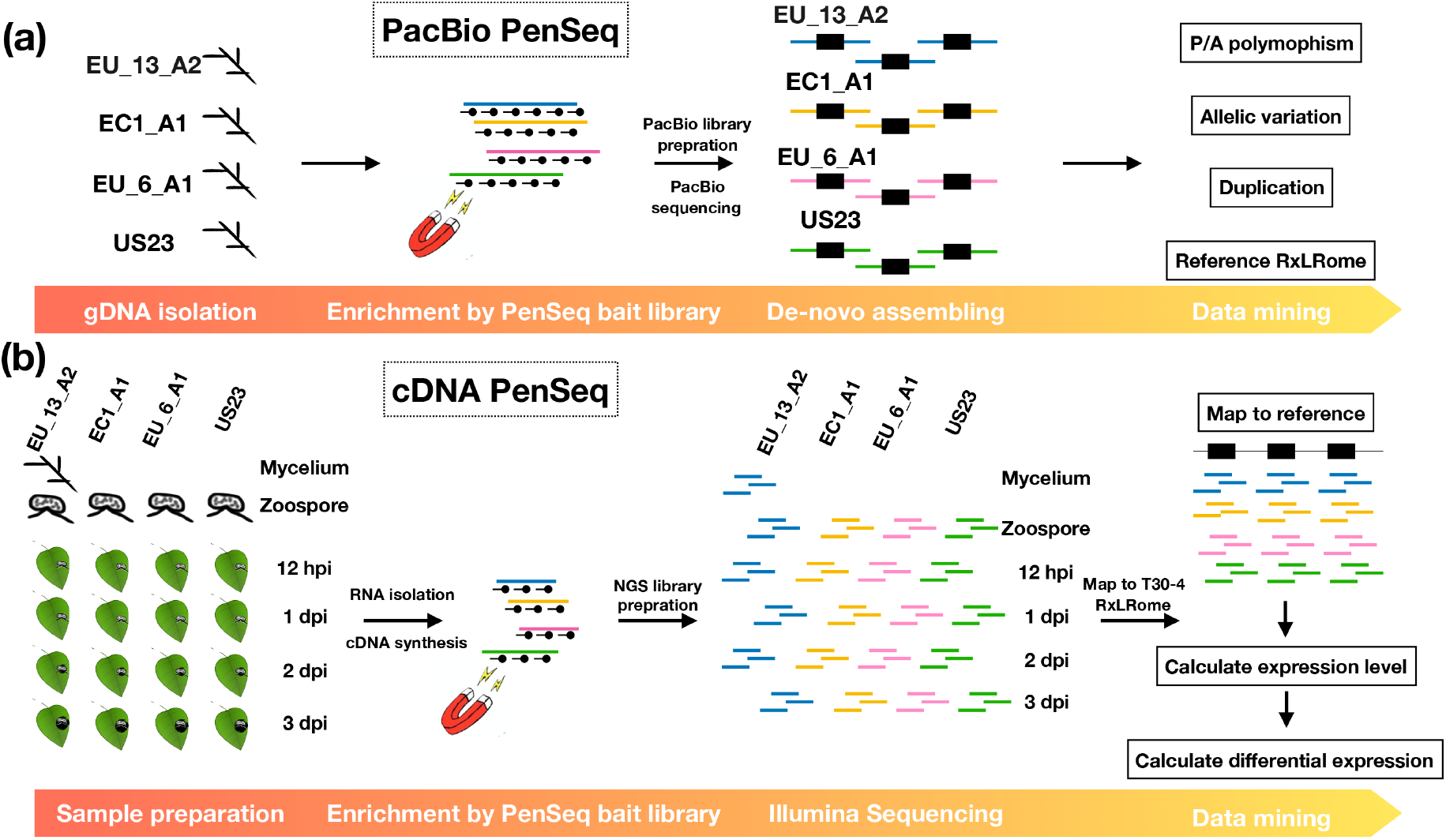
The pipelines of PacBio and cDNA PenSeq. (a) The pipeline of PacBio gDNA PenSeq. Briefly, the gDNA isolated from various *Phytophthora infestans* was enriched for RxLR effectors, sequenced by PacBio and de-novo assembled for data mining. (b) The pipeline of cDNA PenSeq. The cDNA were synthetized using RNA sampled from various *P. infestans* at different stages (mycelium, zoospore, 12 hpi, 1, 2 and 3 dpi). The libraries enriched for RxLR effectors were sequenced, reads were mapped to the RxLRome of the reference *P. infestans* genome T30-4 and the expression levels of samples were calculated and compared. Black lines with dots represent the baits, the enriched fragments are depicted in blue (EU_13_A2), yellow (EC1_A1), pink (EU_6_A1) and green (US23). The black boxes indicate RxLR effectors. EU_13_A2, EC1_A1, EU_6_A1, US23, *P. infestans* genotypes.

Following the enrichment sequencing, circular consensus sequencing (CCS) reads were assembled (Fig. 1a) and contigs with fewer than 10 reads were removed. The average length of the contigs of coverage >10 reads was 7 kb (Table 1), and the size of the largest contig was over 50 kb. This suggests that the PacBio PenSeq successfully captured the target effector genes and the adjacent flanking DNA sequences. In total, 1,137, 1,054, 1,283 and 925 contigs were obtained from EU_13_A2, EC1_A1, EU_6_A1 and US23 respectively, of which 687, 650, 741 and 571 contigs contain RxLR effectors. (Table 1, Notes S1-S4). The remaining contigs contained non-RxLR effectors which were included in the bait library design for other purposes (Thilliez *et al*., 2018).

**Table 1.**
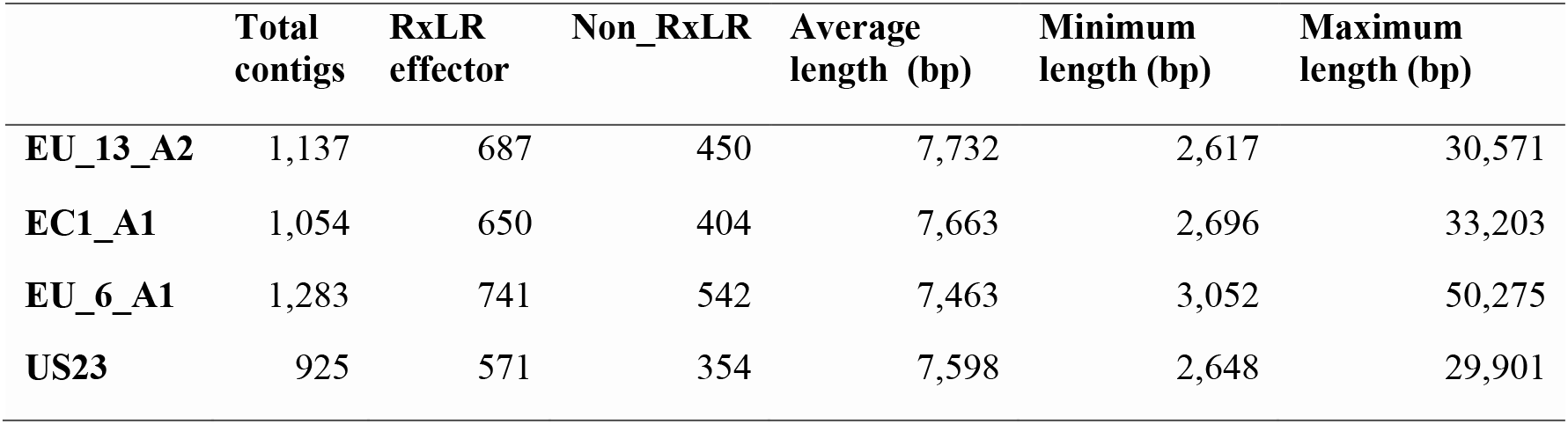
PacBio PenSeq for EU_13_A2, EC1_A1, EU_6_A1 and US23.

The PacBio PenSeq data allowed us to detect new RxLR effector alleles from different haplotypes of various *P. infestans* isolates, and even in polyploid genotypes like EU_13_A2 (Li *et al.*, 2017). This dataset can also be used to extensively study allelic variation, presence/absence (P/A) polymorphism and effector evolution. For example, *Avr1* (PITG_16663) and a paralogous *Avr1-like* gene (PITG_06432) are located on supercontigs 1.51 and 1.8 of the reference T30-4 genome, respectively. The *R1*-breaking clonal lineage EU_13_A2 was reported to have an 18 kb deletion comprising the *Avr1* locus (Cooke *et al.*, 2012). Also, the Illumina PenSeq data showed that the *Avr1* locus is missing in EU_13_A2, EC1_A1 and US23 (Thilliez *et al.*, 2018). We mapped the four *Avr1* contigs from EU_13_A2 contig 192, 261, 296 and 329) to supercontig 1.51 and 1.8, and found that all four contigs map to the *Avr1-like* supercontig 1.8. Two contigs (contig 261 and 286) mapped to the *Avr1-like* locus, and two other contigs (contig 192 and 329) mapped to a locus next to *Avr1-like* that was not previously annotated (Fig. S2), though the genes in those two contigs might be pseudogenes as the signal peptide is missing in both of them. Additionally, in EU_6_A1 and US23, two *Avr1* contigs did not map to *Avr1* or *Avr1-like* loci of T30-4. Thus, our PacBio PenSeq dataset can provide means to detect novel RxLR effector paralogs absent from the reference genome.

As another example, our dataset carries in total 504 of the 563 predicted RxLR effectors from the reference genome T30-4 (Haas *et al.*, 2009). To investigate P/A polymorphism of RxLR effectors in the four sequenced isolates, we performed a basic local alignment search (BLAST) of the 504 effectors against the PacBio contigs, with hits with < 50% coverage defined as absent Table S2). We found that 17, 28, 15 and 33 RxLR effectors out of the 504 are missing in EU_13_A2, EC1_A1, EU_6_A1 and US23, respectively.

Taken together, we have generated a rich dataset that could help to define full length RxLR effector genes, deliver robust information on alleles and paralogs, and reveal conserved or race-specific effectors from different isolates. It is available in full in Notes S1-S4.

### cDNA PenSeq enables effector expression detection in early stages of infection

To clarify whether the untested effectors might be putative *Avr* genes, we performed cDNA PenSeq for the four *P. infestans* isolates EU_13_A2, EC1_A1, EU_6_A1 and US23, at different time points post-infection (12 hpi, 1, 2 and 3 dpi) and in mycelium and zoospores (Fig. 1b). To analyse and visualize the cDNA PenSeq data, we built an artificial DNA sequence contig (“RxLRome”) for the RxLR effectors. In addition, nine non-RxLR genes from the bait library were included as controls (Jouet et al., 2018; Fig. 2). The cDNA PenSeq reads were mapped to the RxLRome and gene expression compared over time (Fig. 1b).

**Fig. 2.**
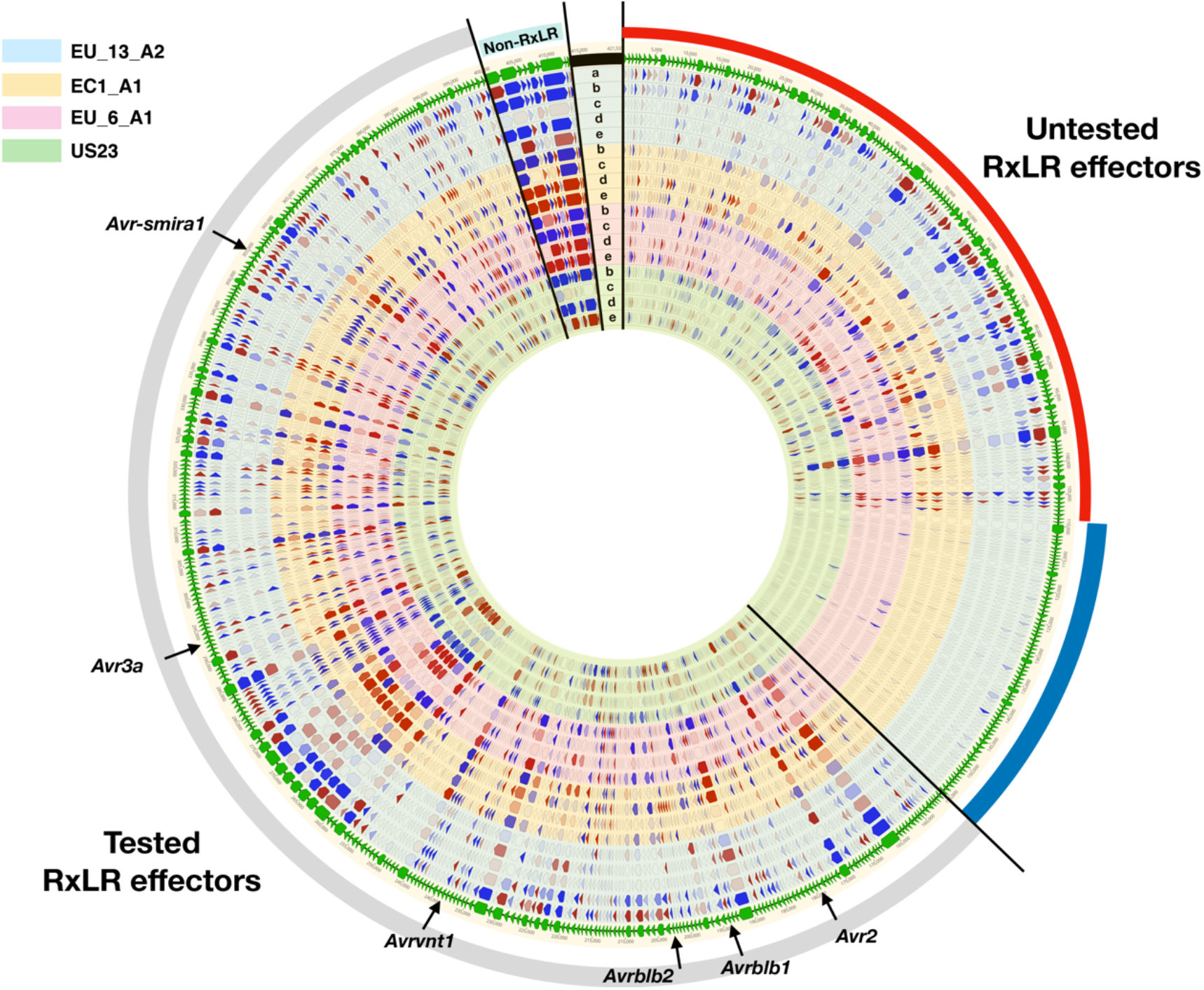
cDNA PenSeq of RxLR effectors from EU_13_A2, EC1_A1, EU_6_A1 and US23. The cDNA PenSeq data for the RxLR effectors from four *Phytophthora infestans* at different stages were mapped to an artificial contig (RxLRome) of 499 RxLR effectors and nine non-RxLR genes, demarcated by bright green arrows on the outer edge of the diagram. Black lines separate the previously tested RxLR effectors (grey bar), new effector candidates with differential expression (red bar), unexpressed effectors (blue bar) and non-RxLR controls (cyan). The concentric circles in blue, yellow, pink and green represent data from *P. infestans* EU_13_A2, EC1_A1, EU_6_A1 and US23, respectively. The arrows on them indicate differential expression (red, up-regulation, blue, down-regulation; no fill, no difference), where the more intense the colour, the bigger the difference. The data are plotted as follows: a, mycelium vs zoospores (for EU_13_A2 only); b, zoospores vs 12 hpi; c, 12 hpi vs 1 dpi; d, 1 dpi vs 2 dpi; e, 2 dpi vs 3 dpi. Six known *Avr* genes, *AvrSmira1*, *Avr3a*, *Avrvnt1*, *Avrblb2*, *Avrblb1* and *Avr2* are indicated by black arrows.

Most of the RxLR effectors which were included in previous effector libraries show an up-regulation of expression in the early stages of infection (Fig. 2). Some of the untested RxLR effectors show a similar pattern of expression, and might also represent potential *Avr* genes, while others are poorly expressed in some isolates. The details of the cDNA PenSeq are available in Table S3.

### Identification of *Avramr1*

To test if *Avramr1* is among the untested effectors, we selected 47 highly expressed effectors (Fig. 3) present in all tested lineages that had not previously been investigated. The effectors were synthesized, cloned into an expression vector with 35S promoter and transformed into *Agrobacterium* GV3101 for agro-infiltration in *N. benthamiana* (Fig. 4a). All the effectors were infiltrated alone or co-infiltrated with *Rpi-amr1-2273* (Witek *et al.*, 2020). Among the 47 effectors, PITG_07569 was the only effector which triggered an HR when co-expressed with *Rpi-amr1-2273* (Fig. 4b). Hence, we concluded PITG_07569 is *Avramr1*.

**Fig. 3.**
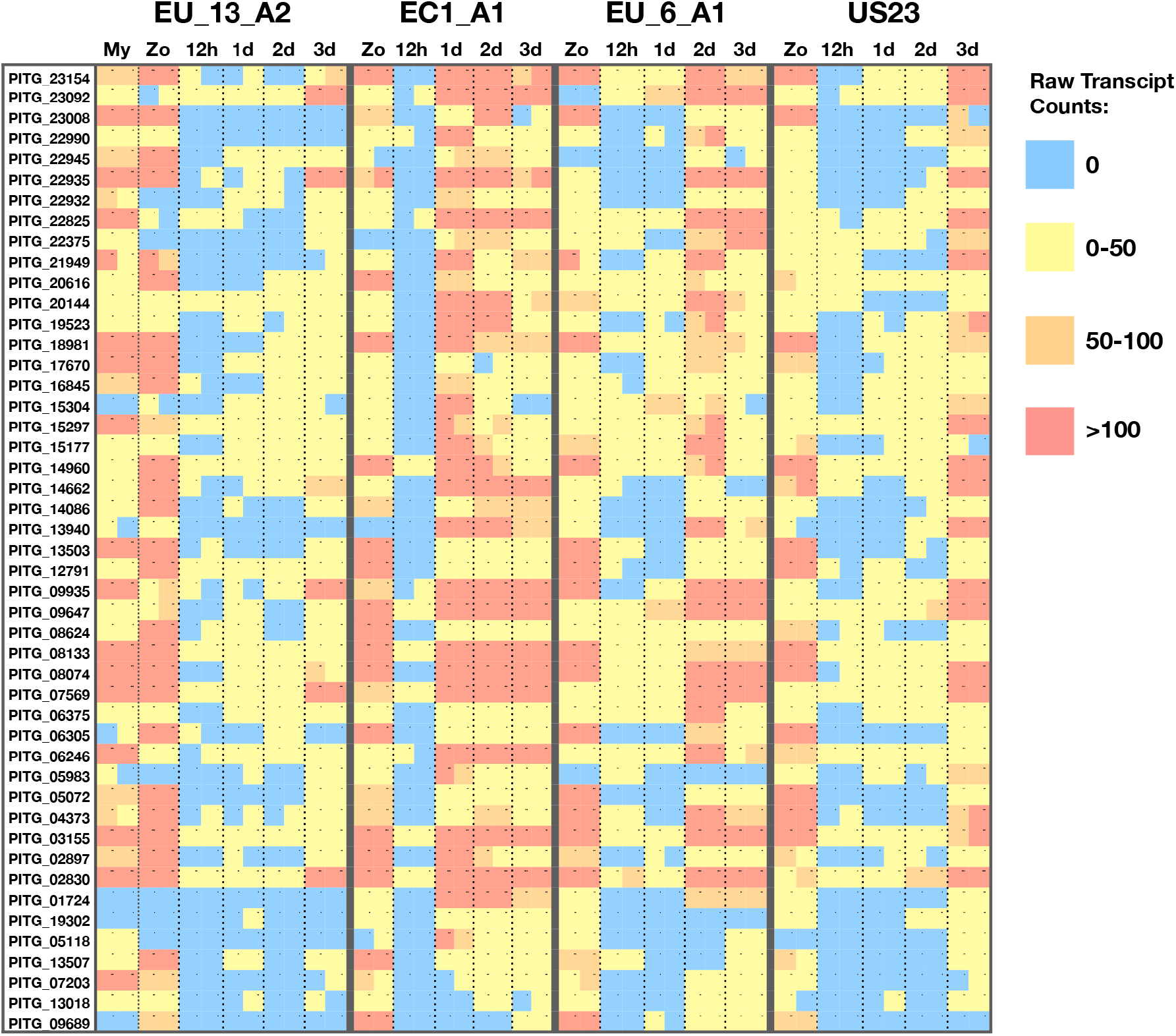
Raw transcript counts for the new candidate RxLR effectors. 47 most differentially expressed RxLR effectors from the previously untested set were selected, and the raw transcript counts were visualized as a heat map across time points and treatments. Each square indicates a single data point derived from two independent biological replicates. The colours red, orange, yellow and blue represent >100, 50-100, 0-50, or 0 raw transcripts, respectively.

**Fig. 4.**
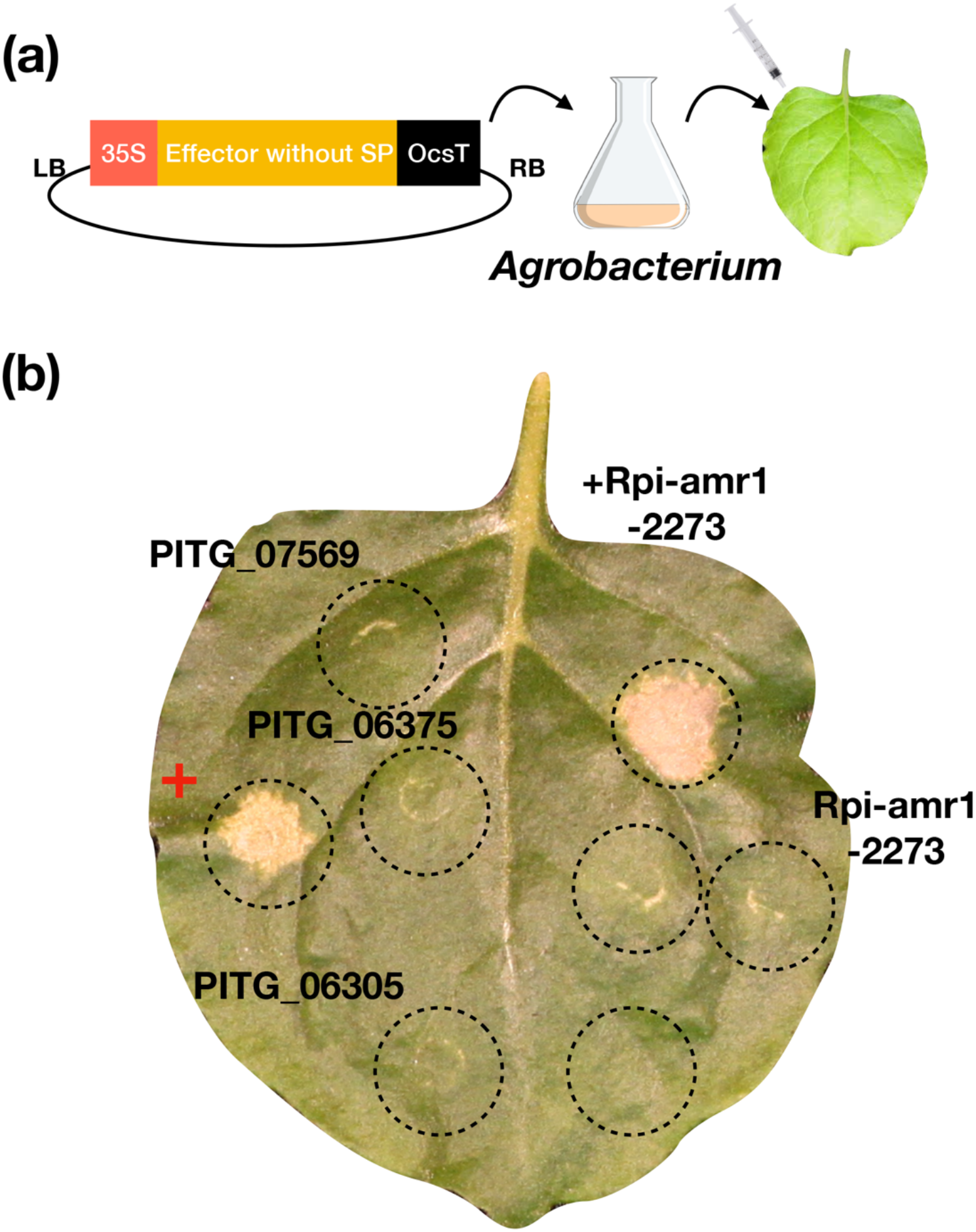
Identification of *Avramr1*. (a) All 47 selected effectors without signal peptides (SP) were synthesized and cloned into an expression vector under CaMV-35S promoter for *Agrobacterium*-mediated transient expression. (b) Transient expression of candidate effectors on their own or with *Rpi-amr1-2273* in *Nicotiana benthamiana*. Dashed circles demarcate the infiltration sites. Only PITG_07569 triggers HR when co-expressed with *Rpi-amr1-2273*. All other effectors that did not trigger HR are represented by PITG_06375 and PITG_06305. A known *R*/*Avr* gene pair was used as positive control (+). This experiment was repeated more than 10 times with the same results.

### *Avramr1* homologs in different *P. infestans* isolates and other *Phytophthora* species

*Avramr1* is a canonical RxLR effector with RYLR and EER motifs and an N-terminal signal peptide (Fig. 5b). *Avramr1* locates on supercontig 1.11 of the *P. infestans* reference genome T30-4. *Avramr1-like* (hereafter *Avramr1L*), a truncated paralog (PITG_07566) maps adjacent to *Avramr1* (Fig. 5a and 5b). Two known *Avr* effectors, *Avr8* (PITG_07558) and *Avrsmira1* (PITG_07550), are physically close to the *Avramr1* locus in the T30-4 genome (Fig. 5a) (Rietman *et al.*, 2012).

**Fig. 5.**
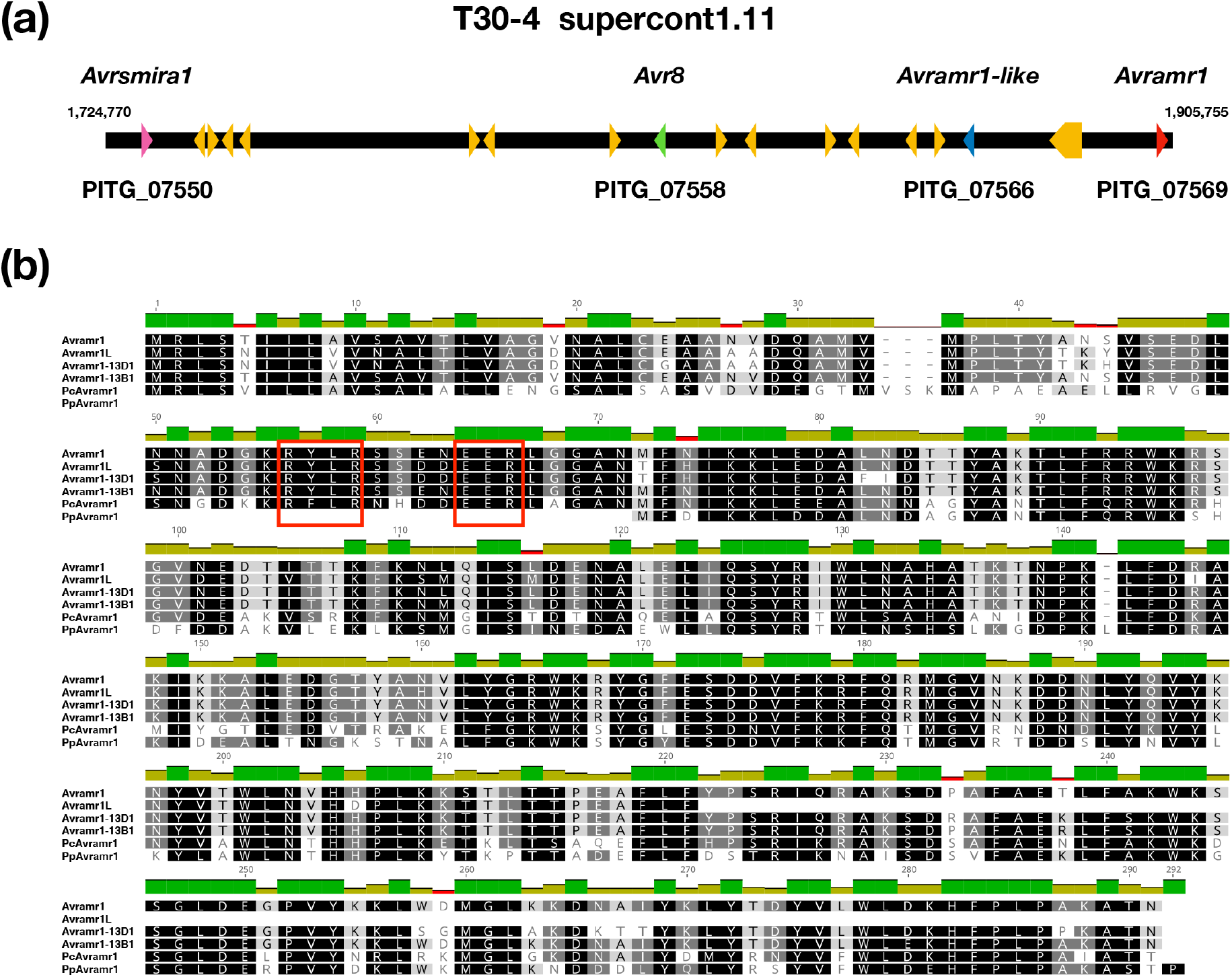
Genomic localization and amino-acid alignment of *Avramr1*. (a) The localization of *Avramr1* (*PITG_07569*, red arrow) on supercontig 1.11 of the reference *Phytophthora infestans* T30-4 genome. A paralog *Avramr1L* gene (*PITG_07566*, blue arrow) is located close to *Avramr1*. The supercontig contains another two known *Avr* genes, *Avrsmira1* (*PITG_07550*, pink arrow) and *Avr8* (*PITG_07558*, green arrow). (b) The alignment of protein sequences of Avramr1 and selected homologs and paralogs from *P. infestans*, *P. capsici* (Pc) and *P. palmivora* (Pp). The dark green bars on top of the alignment indicate 100 % identity while olive green and red bars indicate various degrees of polymorphism between the sequences. RxLR and EER motifs are highlighted by red boxes.

To study the sequence polymorphism of *Avramr1* homologs in *P. infestans*, we used BLAST to search for *Avramr1* homologs in the PacBio PenSeq assemblies generated in this study. It revealed that EU_13_A2, EC1_A1, EU_6_A1 and US23 carry six, four, three and six *Avramr1* homologs, respectively. Next, we aligned the corresponding Avramr1 amino acid sequences and generated a neighbourhood joining (NJ) tree for phylogenetic analysis (Fig. 6a). Two *Avramr1* homologs from *Phytophthora parasitica* and *Phytophthora cactorum* were identified from public database, and they were used as an outgroup (Fig. 5b and 6a). Based on the phylogenetic tree, we distinguished four Avramr1 clades, clade A (containing Avramr1 from T30-4) and clade C (with Avramr1L from T30-4), and two more clades, B and D (Fig. 5a). For a more detailed analysis, we selected one *Avramr1* homolog from clade B and one from D (*Avramr1-13B1* and *Avramr1-13D1* from EU_13_A2) and aligned them with *Avramr1* homologs from clade A and C, and with *P. parasitica* and *P. cactorum* homologs. Significant sequence polymorphisms are observed between effectors from different clades (Fig. 5b). Meanwhile, the *Avramr1* homologs within the same clade are almost identical (Fig. 6a).

**Fig. 6.**
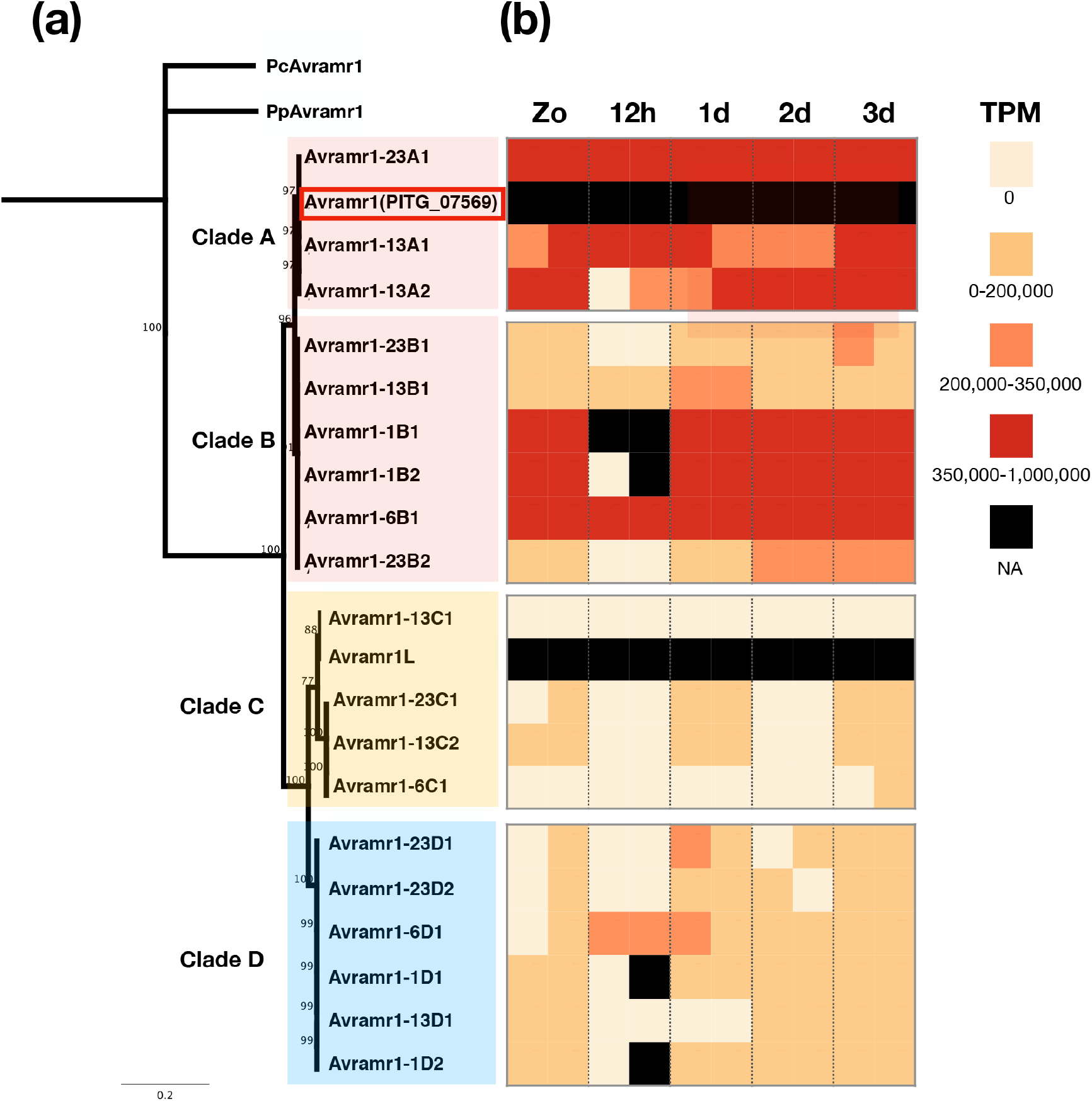
Phylogeny and expression profile of *Avramr1* homologs from EU_13_A2, EC1_A1, EU_6_A1 and US23. (a) Neighbor joining tree of the protein sequences of the *Avramr1* homologs. Pc-Avramr1 and Pp-Avramr1 were used as outgroups. (b) The expression profile of *Avramr1* homologs at different stages and time points (zoospores, 12hpi, 1, 2 and 3 dpi). Transcripts per kilobase million (TPM) for each effector homologs were visualized as follows: black, data not available; red, 350,000-1,000,000 TPM; orange, 200,000-350,000 TPM; yellow, 0-200,000 TPM; beige, 0 TPM.

### Differential expression of *Avramr1* homologs in different *P. infestans* isolates

To investigate the expression patterns of *Avramr1* homologs defined in the PacBio PenSeq data, we mapped the corresponding cDNA PenSeq reads to the PacBio PenSeq contigs from EU_13_A2, EC1_A1, EU_6_A1 and US23. The transcript per million (TPM) values for each time point are visualized in Fig. 6b. The clade A homologs *Avramr1-23A1*, *Avramr1-13A1* and *Avramr1-13A2* are highly expressed at almost all stages, and the *Avramr1* homologs from clade B show a similar expression pattern. For clade C, some homologs, like *Avramr1-13C*, *Avramr1-6C1* or *Avramr1L* gene from T30-4, are weakly expressed at all stages. However, two other *Avramr1L* homologs, *Avramr1-23C1* and *Avramr1-13C2*, show moderately elevated expression in zoospores, and at 1 dpi and 3 dpi. Interestingly, the *Avramr1* homologs from clade D, which are missing in the reference genome T30-4, show an intermediate expression level compared to Clade A, B and Clade C, and most *Avramr1* homologs in Clade D show an increase in expression at the zoospore stage, and at 1, 2, 3 dpi.

In summary, our PacBio PenSeq analysis created a rich dataset to reveal new *Avr* variants from different *P. infestans* isolates, and to quantify their expression profile individually. This facilitates the analysis of the polymorphism of pathogen effectors and their potential differential recognition patterns with the corresponding *Rpi* genes (Witek *et al.*, 2020).

## Discussion

The availability of the *P. infestans* genome sequence enabled a step-change in the rate of investigation of this pathogen, accelerating the discovery of recognized effectors, and of new *Rpi* genes (Haas et al., 2009; Vleeshouwers et al., 2008; 2011). However, some questions remain open. For example, how different are the effector repertoires in different *P. infestans* isolates? To what extent do they show differential expression between races? The study of plant *NLR* gene repertoires faces similar challenges, and sequence capture, combined with long-read sequencing technologies, has enabled the refinement of tools to cost-effectively investigate diversity, such as RenSeq, SMRT RenSeq, RLP/KSeq and AgRenSeq (Arora et al., 2019; Lin *et al.*, 2020; Jupe et al., 2013; Witek et al., 2016). Recently, the pan-NLRome of 65 diverse *Arabidopsis thaliana* accessions was determined by a similar strategy, revealing that any one accession lacks many of the NLRs found in the species pan-NLRome (Van de Weyer *et al.*, 2019).

Pathogen-enrichment sequencing (PenSeq) was developed to facilitate cost-effective investigation of pathogen diversity on infected plants, and polymorphism of pathogen effectors (Jouet et al., 2018; Thilliez et al., 2018). The first PenSeq studies, however, were conducted using Illumina short reads. This significantly limited their resolving power as many oomycete genomes are highly heterozygous, and some effectors belong to large gene families with multiple sequence-related paralogs that can lead to false assemblies (Gilroy et al., 2011; Oliva et al., 2015).

In this study, we combined long read PenSeq and cDNA Penseq, enabling a detailed analysis of the RxLR genes and their expression patterns in different *P. infestans* isolates. The cDNA PenSeq dataset allowed us to define an additional set of 47 RxLR genes expressed during infection that were not previously investigated. Amongst these, we identified *Avramr1*, which encodes the cognate recognized effector for *Rpi*-*amr1* from *S. americanum* (Witek *et al.*, 2020).

The long read PenSeq data helped us to obtain full-length RxLR effector haplotypes with their flanking sequences. This allowed us to distinguish individual alleles from polyploid isolates like EU_13_A2, and also distinct effector paralogs. The sequences flanking the *RxLR* genes enabled us to understand the possible translocation events and identify new *RxLR* loci. We were also able to identify multiple new *Avramr1* homologs from different isolates, and identified a new *Avramr1* Clade D which is not present in T30-4. The PenSeq dataset constitutes a valuable community resource for investigating the allelic and expression diversity of multiple recognized effectors.

So far, no *Rpi-amr1*-breaking *P. infestans* isolates have been found (Witek *et al.*, 2020), and therefore we propose that *Avramr1* might be crucial for the virulence of *P. infestans*. The identification of *Avramr1* will enable us to study its virulence function, its polymorphism in the *P. infestans* population and its recognition by *Rpi-amr1*. Collectively, these data and methods will contribute to understanding this fast-evolving and destructive oomycete pathogen, and to achieving durable late blight resistance in potato.

## Acknowledgements

This research was finance from BBSRC grants BB/P021646/1. We would like to thank Sophien Kamoun and Joe Win for valuable discussion, and TSL Bioinformatics Team, SynBio Team and horticultural team for their support.

## Supporting Information

**Notes S1:** PacBio PenSeq contigs of EU_13_A2.

**Notes S2:** PacBio PenSeq contigs of EC1_A1.

**Notes S3:** PacBio PenSeq contigs of EU_6_A1.

**Notes S4:** PacBio PenSeq contigs of US23.

**Table S1:** 278 RxLR effectors from previously available effector libraries.

**Table S2:** P/A polymorphism of RxLR effectors from EU_13_A2, EC1_A1, EU_6_A1 and US23.

**Table S3: cDNA PenSeq for** EU_13_A2, EC1_A1, EU_6_A1 and US23.

**Fig. S1:** Enrichment efficiency with/ without non-adaptor-ligated DNA.

**Fig. S2:** Compare EU_13_A2 *Avr1* contigs and the T30-4 reference genome.

