## Supplementary material for "Identification of *Avramr1* from *Phytophthora infestans* using long read and cDNA pathogen-enrichment sequencing (PenSeq)": Fig. S1

**With non-adaptor-ligated *P. infestans* DNA**

| | | Before | Post | -ΔCt |
|---|---|---|---|---|
| Length | Gene ID | Mean Ct | Mean Ct | Before - Post |
| 252 | PITG_07482 | 24.05484125 | 17.86351082 | 6.191330427 |
| 285 | PITG_06092 | 23.6159711 | 17.47112222 | 6.144848877 |
| 321 | PITG_04097 | 24.61569113 | 18.90343592 | 5.712255209 |
| 1200 | PITG_10540 | 23.98549956 | 16.45730254 | 7.528197019 |
| 1452 | PITG_06478 | 23.6432591 | 16.77172378 | 6.871535316 |
| 2052 | PITG_05014 | 24.25946711 | 16.22825301 | 8.031214095 |
| 2082 | PITG_15105 | 24.95389085 | 16.09825722 | 8.855633628 |
| 393 | PITG_02116 | 23.65059279 | 29.09760449 | -5.447011704 |
| 891 | PITG_01775 | 23.1732859 | 33.75774971 | -10.58446381 |
| 1140 | PITG_06710 | 23.78762221 | 29.68986618 | -5.902243964 |
| 1068 | PITG_17354 | 24.0055913 | 39.72802676 | -15.72243546 |

**Without non-adaptor-ligated *P. infestans* DNA**

| | | Before | Post | -ΔCt |
|---|---|---|---|---|
| Length | Gene ID | Mean Ct | Mean Ct | Before - Post |
| 252 | PITG_07482 | 23.88495129 | 19.0243649 | 4.860586393 |
| 285 | PITG_06092 | 23.04466229 | 18.66210318 | 4.382559117 |
| 321 | PITG_04097 | 24.3443166 | 19.69343242 | 4.650884185 |
| 1200 | PITG_10540 | 23.44467276 | 17.69033439 | 5.754338375 |
| 1452 | PITG_06478 | 23.4029303 | 17.807447 | 5.595483301 |
| 2052 | PITG_05014 | 23.94112982 | 17.5258043 | 6.415325527 |
| 2082 | PITG_15105 | 24.53579402 | 17.48475851 | 7.051035516 |
| 393 | PITG_02116 | 23.40810244 | 27.5390805 | -4.130978056 |
| 891 | PITG_01775 | 22.91475665 | NaN | NaN |
| 1140 | PITG_06710 | 23.43683537 | NaN | NaN |
| 1068 | PITG_17354 | 23.45621888 | NaN | NaN |
