## Supplementary figures and images for "Identification of *Avramr1* from *Phytophthora infestans* using long read and cDNA pathogen-enrichment sequencing (PenSeq)"

### Fig. S2

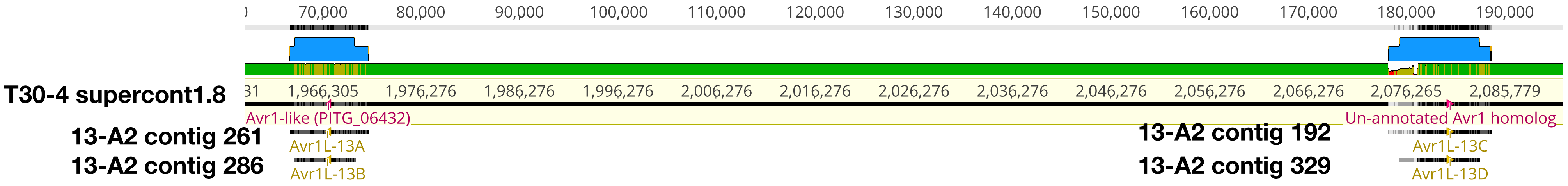
